# Microbial network, phylogenetic diversity and community membership in the active layer across a permafrost thaw gradient

**DOI:** 10.1101/143578

**Authors:** Rhiannon Mondav, Carmody K McCalley, Suzanne B Hodgkins, Steve Frolking, Scott R Saleska, Virginia I Rich, Jeff P Chanton, Patrick M Crill

**Author notes:** Corresponding author: RMondav, Norbyvagen, Uppsala University, Uppsala, SE75236 Sweden. Current addresses: Thomas H. Gosnell School of Life Sciences, Rochester Institute of Technology, Rochester, New York 14623, USA. Department of Microbiology, The Ohio State University, Columbus 43210, USA.

## Abstract

Biogenic production and release of methane (CH_4_) from thawing permafrost has the potential to be a strong source of radiative forcing. We investigated changes in the active layer microbial community of three sites representative of distinct permafrost thaw stages at a palsa mire in northern Sweden. The palsa sites with intact permafrost, and low radiative forcing signature had a phylogenetically clustered community dominated by *Acidobacteria* and *Proteobacteria.* The bog with thawing permafrost and low radiative forcing signature was dominated by hydrogenotrophic methanogens and *Acidobacteria*, had lower alpha diversity, and midrange phylogenetic clustering, characteristic of ecosystem disturbance affecting habitat filtering, shifting from palsa-like to fen-like at the waterline. The fen had no underlying permafrost, and the highest alpha, beta and phylogenetic diversity, was dominated by *Proteobacteria* and *Euryarchaeota,* and was significantly enriched in methanogens. The mire microbial network was modular with module cores consisting of clusters of *Acidobacteria, Euryarchaeota,* or *Xanthomonodales.* Loss of underlying permafrost with associated hydrological shifts correlated to changes in microbial composition, alpha, beta, and phylogenetic diversity associated with a higher radiative forcing signature. These results support the complex role of microbial interactions in mediating carbon budget changes and climate feedback in response to climate forcing.

## Introduction

Modern discontinuous permafrost is found in regions with a mean annual air temperature between -5 °C and +2 °C and where the insulating properties of peat enable persistence of permafrost above freezing temperatures (Shur and Jorgenson, 2007; Seppälä, 2011). As these regions experience climate change-induced warming, they are approaching temperatures that are destabilizing permafrost (Schuur et al., 2015). Permafrost degradation typically leads to significant loss of soil carbon (C) through erosion, fire and microbial mineralisation (Osterkamp et al., 2009; Mack et al., 2011). The area of permafrost at risk of thaw in the next century has been estimated to be between 10^6^ and 10^7^ km^2^, with the quantity of C potentially lost ranging from 1-4 x 10^14^ kg (McGuire et al., 2010; Schuur et al., 2015). Increasing plant production in thawed systems may partially compensate for this loss but this is poorly defined (Hicks Pries et al., 2013). Thus, the potential positive feedbacks to climate change are not well constrained, and will vary depending on the emission ratio of the greenhouse gases (GHGs) carbon dioxide (CO_2_) and methane (CH_4_), and the time scale considered (Dorrepaal et al., 2009; Nazaries et al., 2013). Changes in microbial community membership (e.g. methanogen to methanotroph ratio) will be a significant factor controlling the CO_2_ to CH_4_ emission ratio (Hodgkins et al., 2014).

To examine the relationship between permafrost thaw, shifting GHG emissions, and the microbial community, we investigated a natural *in situ* thaw gradient at Stordalen Mire, northern Sweden, on the margin of the discontinuous permafrost zone. Permafrost thaw has been causatively linked to changes in topography, vegetation, and GHG emission at Stordalen (Christensen, 2004; Johansson et al., 2006; Bäckstrand et al., 2008, 2010; Johansson and Åkerman, 2008), and elsewhere (Turetsky et al., 2007; Olefeldt et al., 2013). Currently, the Mire is a partially degraded complex of elevated, drained hummocks (palsas with intact permafrost) and wet depressions (bogs with thinning permafrost and fens without any permanently frozen ground), each representing different stages of thaw (Fig S1), and each characterised by distinct vegetation (Bhiry and Robert, 2006; Johansson et al., 2006). Changes in topography and vegetation (proxy for thaw) have been tracked through the last 40 years and show a decrease in area covered by palsa, an expansion of fens and a variable area covered by bogs (Christensen et al., 2004; Malmer et al., 2005; Johansson et al., 2006). Part of the known palsa-cycle is the external carving and internal collapse of palsas into the surrounding bog or fen, though the time scale at which this happens varies depending on whether dome-palsas or palsa-complexes/plateaus (as seen at Stordalen) are considered (Railton and Sparling, 1973; Zoltai, 1993; Sollid and Sørbel, 1998; Gurney, 2001; Turetsky et al., 2007; Seppälä, 2011; O’Donnell et al., 2012; Liebner et al., 2015). At Stordalen complete collapse due to absence of permafrost results in a fen or lake, while partial collapse due to permafrost thinning results in a bog (Johansson et al., 2006). Photographs, topographical survey and GHG data comparing Stordalen in the 1970’s and 1980’s to 2000’s and 2010’s show that for the particular area studied here the palsa has degraded both externally and internally (Fig S1, S2). Bogs (sphagnum or semi-wet) have expanded within the perimeter of the palsa-complex and around its southern edge, while fens (eriophorum, wet, or tall-graminoid) have encroached from the north, east, and west having converted the bog that once existed on the western and eastern edges of the palsa, along with increases in GHG emissions (Rydén et al., 1980; Malmer et al., 2005; Johansson et al., 2006; Bäckstrand et al., 2010). The majority of bog samples were taken from the subsided section within the palsa-complex, while the fen samples were taken from the western side of the complex, which was recorded as a bog 40 years ago. As permafrost continues to disappear from Stordalen over the coming decades, subsidence of the surface will likely increase, driving the creation of more transient bog-type communities and degradation into fens (Christensen, 2004; Parviainen and Luoto, 2007; Johansson and Åkerman, 2008; Fronzek et al., 2010; Bosiö et al., 2012; Jones et al., 2016). It is also predicted that this region could be free of permafrost as early as 2050 (Parviainen and Luoto, 2007; Fronzek et al., 2010). The “natural experiment” underway in the Mire presents a model ecosystem for investigating climate-driven changes in lowland permafrost regions with high cryosequestered-C (Masing et al., 2009).

As a model system, the Mire has been intensively studied over the last several decades for permafrost thaw impacts on plant communities, hydrology, and biogeochemistry – providing rich context for interpreting microbial communities. The seasonally thawed peat layer (active layer) of Stordalen’s palsas is drained, aerobic, ombrotrophic (rain-fed) and isolated from nutrient-rich groundwater. The palsa sites’ low plant productivity and aerobic decomposition make them net emitters of appreciable CO_2_ and no CH_4_, with a net C balance (NCB) of - 30 mgC/m^2^/day (negative value indicates net C emissions, positive indicates net uptake; Bäckstrand et al., 2010). In contrast, the bog sites (semi-wet in Johansson *et al* 2006) are physically lower and collect rainwater, leading to partial inundation, and are dominated by layered bryophytes (typically *Sphagnum* spp.). Although this results in less lignin (a recalcitrant C compound not produced by sphagnum), C degradation is still slow due to sphagnum’s higher phenolic content (Freeman et al., 2004) and extremely poor C:N ratios of up to 70:1 (Hodgkins et al., 2014). Partially anoxic conditions permit microbial fermentation and CH_4_ production (Nilsson and Bohlin, 1993; Hobbie et al., 2000). Mire bog sites have the lowest radiative forcing signature (NCB in CO_2_ eq. of - 8 mgC/m^2^/day; Bäckstrand et al., 2010), as fixation of C in the bog peat is high compared to emission of C gases. Finally, the fully-thawed fen sites (tall-graminoid in Johansson *et al* 2006) are the most subsided and are minerotrophic (groundwater-fed). Vegetation succession results in dominance of graminoids (sedges, rushes, reeds), with a subsequent shift in the litter preserved as peat. Some graminoids enhance gas transport between inundated soil and the atmosphere (Chanton et al., 1993) and due to high productivity, contribute appreciable fresh labile organic litter and exudates (Wagner and Liebner, 2009). High productivity results in the fens being the Mire’s biggest gross C-sinks of the Mire, however their high CH_4_ emissions result in a net warming potential 7 and 26 times greater than the palsa and bog respectively (NCB in CO_2_ eq. of - 213 mgC/m^2^/day, Bäckstrand et al., 2010; Christensen et al., 2012).

Here we explore the relationship between the biogeochemical differences among palsa, bog, and fen and the active layer microbial community, via a temporal (over a growing season) and spatial (across habitats, and with depth through the active layer) community survey using SSU rRNA gene amplicon sequencing, pore-water chemistry, peat chemistry, stable C isotope analyses, and CH_4_ flux. Previous work has demonstrated that permafrost thaw has an overall impact on Stordalen Mire’s microbiota (Mondav & Woodcroft et al, 2014; Hodgkins et al., 2014; McCalley et al., 2014). Here, we deepen understanding of this impact by addressing the following descriptors with respect to climate-induced thaw and correlated environmental parameters: 1) dominant phyla, 2) beta diversity, 3) assemblage alpha diversity, 4) phylogenetic distance to identify drivers of assembly processes, 5) community network, and 6) C-cycling phylotype distribution.

## Results and discussion

### Dominant phyla of each thaw stage

Dominant phyla can inform on geochemical correlations and community functionality, reflecting overall habitat conditions. *Acidobacteria* and *Proteobacteria* were ubiquitous phyla across the Mire (Fig 1). Dominant palsa phyla also included *Actinobacteria* and Candidate bacterial phylum “WD272” (WPS-2), the abundances of which decreased across the thaw gradient (palsa>bog>fen). Surface bog samples had similar community composition to palsa samples being dominated by *Acidobacteria* and *Proteobacteria* suggesting that site classification, which was identified by vegetation, is not the only environmental correlate important to microbial community assembly in the Mire. Bog samples at or below the waterline (midpoint and deepest) retained similar proportions of *Acidobacteria* and *Actinobacteria* as palsa samples. *Proteobacteria* however, were relatively less abundant, likely due to lower C lability (Goberna et al., 2014; Hodgkins et al., 2014), and were replaced by *Euryarchaeota* in deeper anoxic samples. The shift in phylum ratios from palsa-like to fen-like supports the transitional nature of this thawing site. The most abundant phyla in the fen were *Euryarchaeota, Proteobacteria, Bacteroidetes,* and *Chloroflexi* followed by *Acidobacteria. Woesearchaeota* (DHVEG-6) were only detected in fen samples.

**Fig 1.**
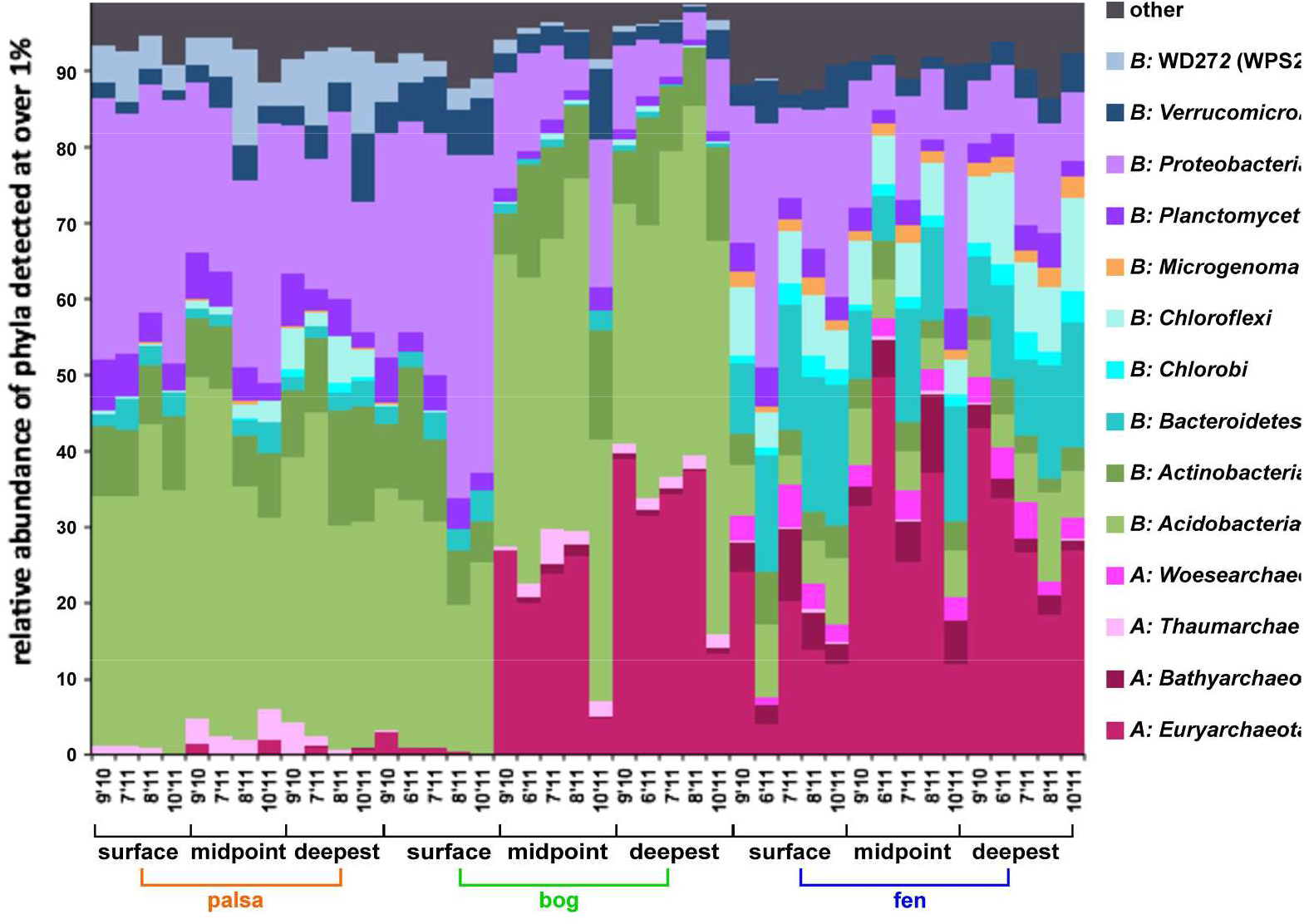
Mean relative abundance of phyla present at over 1 %. Samples grouped by site, then depth, then date. Date is marked along the horizontal axis as m’yy. 210x148mm (300 x 300 DPI)

## <Fig 1, 110 mm wide>

### Beta diversity and environmental correlates of site assemblages

Microbial assemblages were significantly different (anosim 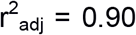, p< 0.001) at the sample OTU level between sites. Whole community analysis by nonmetric multidimensional scaling (NMDS) (Fig 2a, stress = 0.082, r^2^ = 0.99) clustered samples by site. Samples separated along the primary NMDS axis according to hydrological states with ombrotrophic (palsa and bog) samples clustered together left of the origin and the minerotrophic fen samples to the far right. The secondary axis separated samples according to depth from surface with the two ombrotrophic site samples diverging from each other with depth. The palsa and surface bog assemblages overlapped at both OTU (Fig 2) and phyla level (Fig 1). Sharing of species across the palsa and surface bog (aerobic ombrotrophic) is likely due to local dispersal and seen in other methanogenic soils (Kim and Liesack, 2015). Local dispersal mechanisms include transport by burrowing lemmings, palsa dome runoff washing microbes into lower altitudes, local aerial dispersal (Bowers et al., 2011). Ubiquitous deposition across the mire via precipitation may also contribute (Christner et al., 2008). These more ubiquitous microbes likely persist through environmental filtering including oxygenation, acidity, ombrotrophy, and bryophyte presence (Brettar et al., 2011; King et al., 2012).

**Fig 2.**
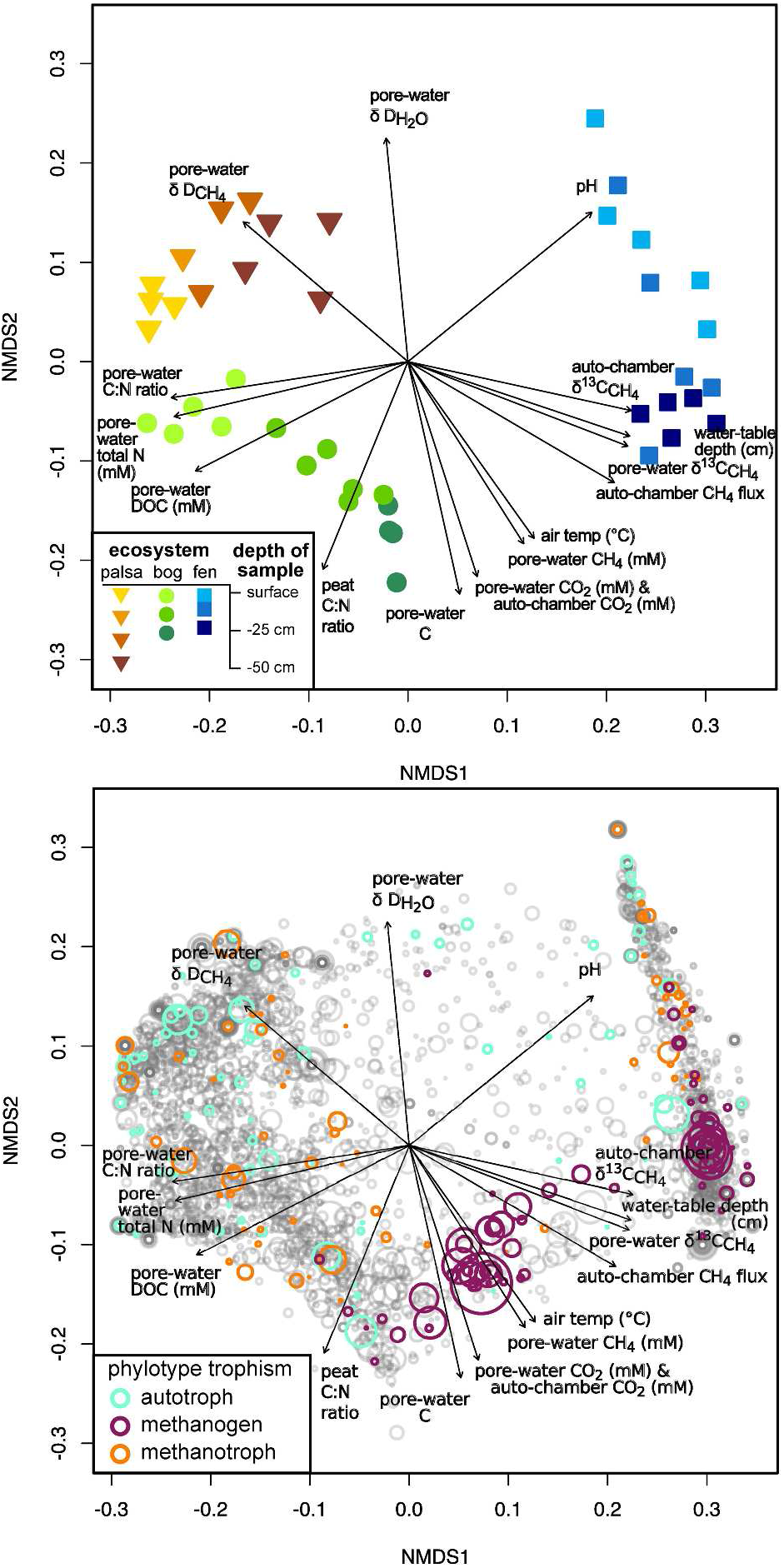
NMDS analysis of sample dissimilarity in community phylotype space, and environmental correlates (Stress = 0.0818, r2 = 0.993). a) The clustering of sites based on their community composition; sites are distinguished by shape and colored to show depth of sample; b) The relative positional contribution of phylotypes to this clustering, with phylotypes plotted as circles with diameter scaled to the log of the mean abundance, and colored to show putative C-cycle metabolism. Plotted vectors on a) & b) are measured environmental variables that had significant correlation to differences in assemblage composition (p < 0.01), with the terminal arrow indicating the direction of strongest change without reference to sign (+ or -). 285x572mm (300 x 300 DPI)

C-fixing autotrophs were associated with all samples except the deepest bog and fen. Bacterial methanotrophs, while distributed across the Mire, were not associated with deep fen or deep bog samples, supporting their known preference for aerobic and microaerobic habitats. Both autotrophs and methanotrophs were unique to individual sites with only a few OTUs shared across the surface bog and palsa samples. The majority of methanogenic phylotypes were clustered exclusively with fen samples, though a secondary cluster of methanogens while more tightly associated to deep bog samples were detected across both bog and fen.

Relative increases in δ D_H2O_ and δ D_CH4_ were correlated with palsa sample assemblages (Fig 2a&b). This is indicative of methanotrophy as supported both by the detection of methanotrophic phylotypes but also by the negative CH_4_ flux (uptake of CH_4_ from atmosphere) recorded by the palsa auto-chambers. Increased peat C:N ratio, pore-water C:N, and porewater DOC were correlated with the bog assemblages supporting previous studies finding higher DOC in run-off from ombrotrophic regions of the Mire (Nilsson, 2006; Kokfelt et al., 2010) Pore-water N content decreased in relation to deeper bog samples. Increased pH, CH_4_ flux from the auto-chambers, flux **δ**^13^C_CH4_ from the autochambers, pore-water **δ** ^13^C_CH4_, and water table depth (WTD) were positively correlated with fen samples (Fig 2a). Pore-water CH_4_ concentration, pore-water CO_2_ concentration, CO_2_ gas from peat samples, and pore-water total C increased with deeper (below watertable) bog and fen samples. Increased CH_4_ in porewater and flux measurements were correlated to presence of detected methanogens (Fig 2b) supporting established linkage between detected abundance and metabolic activity of these microbes (Mondav et al., 2014).

### <Fig 2, 110 mm wide>

#### Microbial assemblage alpha diversity

The number of unique OTUs per normalised (N=2000) sample ranged from 309 in the merged anoxic bog samples from august 2011 up to 1226 in the combined surface fen samples from june 2011 (Fig 3), with a cross-Mire mean OTU richness of 721 (s.d. 204 OTUs). The total richness was 9700 across the Mire for the 42 merged and normalised samples. The percentage of OTUs that could not be taxonomically classified either to or below order level was typical, at 40%, supporting that environmental microbes are still appreciably under-characterised. Total, archaeal, and bacterial assemblage alpha diversity as measured by richness (observed OTUs), Fisher alpha (Fisher et al., 1943), Shannon entropy (Shannon and Weaver, 1949), and Heip’s evenness (Heip et al., 1974) varied between thaw stages, with there being a significant difference (p < 0.05) between sites as measured by Kruskal-Wallace test (K-W) (Fig 3). For total assemblage alpha diversity the bog had lowest (richness and fisher alpha), and was significantly lower than the palsa (shannons entropy and heips evenness) K-W post-hoc test for significance (K-Wmc, p < 0.001). The fen had highest archaeal alpha diversity (richness, fisher, and Shannon) while the palsa had most even archaeal assemblage (K-Wmc, p < 0.001). Apart from the exception of archaeal evenness the bog had lower or lowest alpha diversity of the three sites. Archaeal evenness in the bog site covers a wide range from assemblages with high evenness similar to that found in palsa samples but also includes assemblages that were more highly dominated than those found in the fen (Fig 3). Evenness is an important property of methane producing communities where higher evenness of fen assemblies, compared to bog samples, may constitute a feedback mechanism by which higher CH_4_ production is enabled (Galand et al., 2003; Godin et al., 2012). Depth of sample was related to decreases in all alpha diversity metrics (total and bacterial, range: −0.46 < ρ < −0.36, p < 0.01). Bacterial (r^2^_adj_ = 0.92, p<0.001) and total richness (r^2^_adj_ = 0.92, p<0.001) decreased with depth in the bog while Archaeal richness increased (r^2^_adj_ = 0.69, p<0.001) (Fig 3, Equations S1 a-c). Higher DOC correlated to lower richness (total ρ = -0.76, bacterial ρ = -0.70, and archaeal ρ = -0.74: p<0.001) (Fig S3). Bacterial richness was correlated to decreased porewater CO_2_ (ρ = -0.60, p<0.001, Fig S3). Archaeal richness was positively correlated to distance below water-table (ρ = 0.83, p<0.001, Fig S3). The number of singletons observed in each site directly correlated with richness and varied between sites (r^2^ _adj_ = 0.97, p<0.001, Fig S4, Equations S1 d-g).

**Fig 3.**
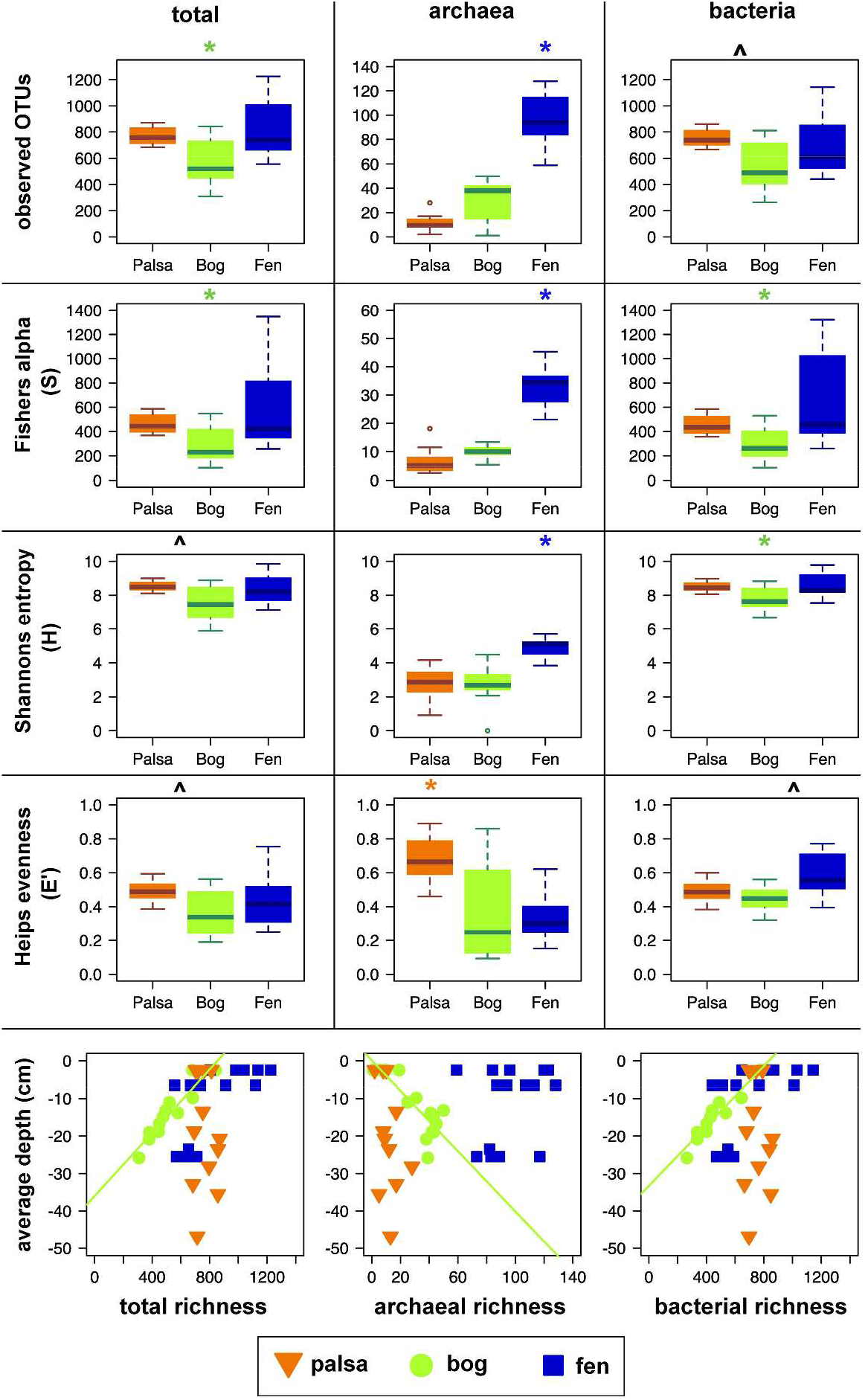
By-site comparison of alpha diversity metrics. a) The distribution of observed richness (top), Fisher’s alpha-S (2nd from top), Shannon’s entropy-H (2nd from bottom), Heip’s evenness (bottom), on all 97 % OTUs (left), archaeal OTUs (middle) and bacterial OTUs (right) in combined normalized (N = 2000) samples. Significant differences were measured by Kruskal-Wallis post-hoc test (* p < 0.001, where * designates a site significantly different from the other two sites and ^ designates a significant difference between the two adjacent sites only)). b) The distribution of OTU richness with sample depth for all OTUs (left), archaeal OTUs (centre), bacterial OTUs (right). Green trend lines show the correlations between richness and depth of sample in the bog site. 301x488mm (300 x 300 DPI)

### <Fig 3, 80 mm wide>

#### Site assembly dynamics

Links have been drawn between a community’s diversity, functional and phylogenetic redundancy (robustness), and the community’s ability to maintain function during change (resistance), and to recover original state if the disturbance is removed (resilience) (Shade et al., 2012; Venail and Vives, 2013). While the degree to which phylogenetic and/or functional diversity influence resistance is not fully elucidated, recent studies support (Werner et al., 2011; Singh et al., 2014), these are important properties to consider with climate change and permafrost thaw, two interacting press disturbances at Stordalen (Shade et al., 2012; Hayes et al., 2014). A press disturbance is a change in the environment that persists for a long period of time, in comparison to a pulse disturbance which is a change that decreases suddenly after a short period of time. Phylogenetic robustness can be measured by various diversity relationships including phylogenetic distance between OTUs (PD); nearest taxon index (NTI), which examines phylogenetic clustering of closely related phylotypes; and mean relatedness index (NRI) which examines variance of phylotype distance within an assemblage. Further, these indices can indicate the relative degree to which stochastic or deterministic processes contribute to community assembly (Wang et al., 2013). The Mire as a whole and each site individually had positive correlation (r^2^ _adj_ = 0.73 to 0.94, p<0.001) between overall phylogenetic diversity (PD) and richness (Eqs S2a-d, Fig S5). Because PD and richness were auto-correlated, subsequent analyses examined PD/OTU so as to examine only the difference due to diversity and not an artefact of abundance counts. Fen PD/OTU was significantly (p<0.001) higher than both bog and palsa assemblages (Fig 4). Measuring assemblage net relatedness (NRI, equivalent to - sesMPD) by OTU phylogenetic distance from sample mean as generated by the null model examines clustering over a whole tree. Greater emphasis is placed on changes towards the tree root compared to other measures such as NTI, which examines diversity at the tips of the phylogenetic tree. Negative values of NRI indicate expansion of the tree via increased branching at higher-level tree nodes i.e. even-dispersal, while positive values indicate filling in of internal phylogenetic tree nodes i.e. clustering. Palsa assemblages had uniformly high NRI, (0.1 < NRI < 0.4, Fig 4) indicative of phylogenetic clustering. A deterministic factor associated with clustering specific to the palsa (and surface bog) is dry-ombrotrophy which increases habitat isolation in heterogeneous soil environments (Kraft et al., 2007; Jones et al., 2009; Stegen et al., 2013; Quiroga et al., 2015). Fen assemblage NRIs were significantly lower than the other sites (KW, p<0.001, Fig 4) and were neutral to negative. Bog assemblage NRIs varied (0 < NRI < 0.4) from neutral in deeper samples to higher than some Palsa in surface samples (Fig 4, Fig S5, Eq S4). The lower NRI in the fen indicates assemblages have broader representation across the bacterial and archaeal domains with less clustering than predicted by the null model i.e. phylogenetic even dispersal. Even dispersal can indicate an assemblage less affected by deterministic processes such as habitat filtering or isolation (Webb et al., 2002; Horner-Devine and Bohannan, 2006), while being more affected by stochastic processes such as dispersal and drift or controversially, by competition (Mayfield and Levine, 2010). All assemblage NTI values were above zero, i.e. more clustered than predicted with the null model, a result seen in early stage successional forests and freshwater mesocosms, (Horner-Devine and Bohannan, 2006; Whitfeld et al., 2012). The bog and fen had greater phylogenetic tip clustering than the palsa (KW p < 0.05, Fig 4) and was, in the bog, related to depth (Eq S3). Greater tip-clustering as seen in the bog and fen indicates greater genomic diversification within ‘species’-populations. This could be achieved through horizontal gene transfer (HGT) or endogenous mutation enabling more closely related organisms to coexist (Goberna et al., 2014), though there is not yet data to address effective population sizes, or the relative frequencies of HGT and endogenous genome mutation in these habitats. The ratio of NRI to NTI indicates the level (tree:tip) at which phylogenetic diversity is affected by assembly processes. Fen assemblages had significantly greater clustering at the whole tree level than tree tip when compared to the other sites (NRI/NTI, Fig 4, p<0.01). Conversely, the palsa assemblages displayed greater clustering towards the tree’s tips. The bog NRI/NTI was in-between the palsa and the fen. This shift in where the assemblage diversity lies supports clustering through local species divergence being a property of the ombrotrophic mire sites while even-dispersal is more prevalent in the minerotrophic fen. The clustering shift from tip to whole tree diversity is also seen at a smaller scale within the bog site (Fig S5, Eq S5) where surface samples have higher tip clustering and deeper samples have more evenly distributed diversity. That the mid depth bog samples were taken at the waterline supports that a main factor regulating this shift is inundation by water. Phylogenetic diversity of Mire assemblages as measured through PD, NRI, and NTI showed the bog grouping alternately with the palsa or the fen supporting that the bog may be an intermediate site undergoing transition from a palsa-like assemblage to fen-like assemblage due to a shift from ombrotrophy to minerotrophy. These four phylogenetic distance analyses support that each site has a unique overall phylogenetic diversity profile, thus giving support to there being differences in assembly and evolution of the microbial community across sites and therefore thaw stages (Stegen et al., 2012).

**Fig 4.**
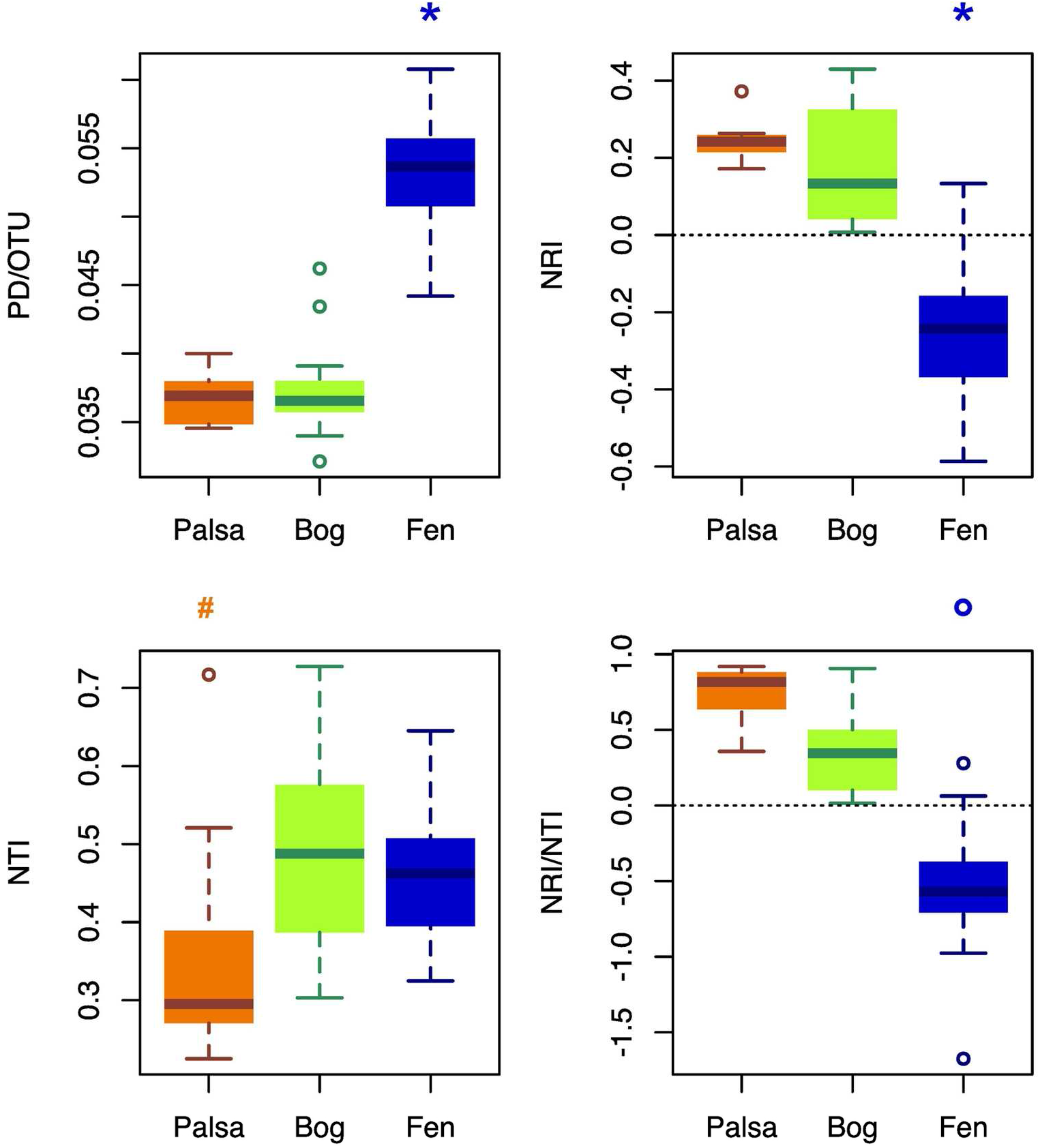
By-site comparison of phylogenetic diversity. topleft: faiths phylogenetic distance per OTU (PD/OTU); bottom left: Nearest taxon index (NTI); top right: Net relatedness Index (NRI), bottom right: NRI/NTI ratio. 147x167mm (300 x 300 DPI)

Examining correlations between environmental parameters and phylogenetic diversity can potentially inform on the relative contribution, directly or indirectly, of environmental to assembly processes. Phylogenetic clustering related to environmental filtering is predicted to be evident in environments with poor nutrient availability or parameters considered to increase selection such as high acidity. Phylogenetic evenness, conversely, is expected to be evident in environments with high resource availability where competition becomes more dominant in assembly processes. Increasing soil pH, ratio of methanogens to methylotrophs, CH_4_ flux, and distance below watertable were significantly correlated to increasing PD/OTU and decreasing NRI and NRI/NTI (ρ ≥ ±0.60, p<0.001, Table S2, Fig S6) i.e. associated with phylogenetic even dispersal. Greater depletion of ^13^C_CH4_, higher porewater DOC, and higher porewater C:N ratios were correlated to decreasing PD/OTU and increasing NRI, and NRI/NTI i.e. phylogenetic clustering (ρ ≥ ±0.60, p<0.001, Table S2). Only bog and fen porewater DOC and C:N were analysed as there was insufficient moisture in the palsa samples. Conditions associated with environmental filtering (acidity) and isolation (ombrotrophy) may here be linked to phylogenetic clustering (Kraft et al., 2007). Stochastic mechanisms specific to the fen include inflow of runoff from the raised palsa and bog, minerotrophy, and local water mixing which reduces isolation (Putkinen et al., 2012). Conditions associated with phylogenetic even-dispersal may therefore be linked to warming potential (increased CH_4_ flux) and increases in acetotrophic methanogens (less depletion of ^13^C in CH_4_ emissions) in methanogenic soils. Clustering is an emerging characteristic of soil communities (Lozupone and Knight, 2007; Auguet et al., 2010) that is currently connected to habitat filtering (Kraft et al., 2007; Shade and Handelsman, 2012) though some recent evidence also supports a role for biotic filtering (Goberna et al., 2014). The correlations between higher phylogenetic diversity, including even-dispersal, and CH_4_ flux corroborate findings from reactors and environmental systems (Werner et al., 2011; Yavitt et al., 2011) that the structure of microbial communities may be significant to global CH_4_ budgets.

### <Fig 4, 80 mm wide>

#### Network topology and community interactions

Microbial networks can be described mathematically by topological indices. Common indices include degree, modularity, betweenness, and closeness. Degree describes the level of connectedness between phylotypes by counting the number of phylotypes that co-occur. Modularity identifies if sub-networks of co-occurrence exist within a larger community network and is thought to be an indicator of resilience. Betweenness Centrality provides information on how critical a phylotype is to the connectedness of a network. Closeness Centrality describes how closely a phylotype is connected to all others in the same module. Redundancy (e.g similar metabolic strategies) is also a useful descriptor of co-existing organisms. Degree, closeness and redundancy in microbial networks provide information on the community’s robustness and, potentially, ability to resist change. OTU Table B_2000_ had 9 700 unique phylotypes, 93% sparsity, average inverse Simpsons (n_eff_) of 122 per sample which was too sparse to obtain meaningful network information from. Restricting the dataset to OTUs that were present in at least 15 samples (one third of total) left 257 unique OTUs, a table with 49% sparsity, and an average n_eff_ of 26, statistics that provide assurance that network interactions could be correctly identified while minimising type I errors (Friedman and Alm, 2012; Berry and Widder, 2014; Weiss et al., 2016). Retaining only OTU pairs that were significantly correlated in at least two of the network analyses from MENA, fastLSA, CoNet (Pearsons, Spearman, Bray, KBL), and SparCC reduced the dataset to 123 OTUs with 265 significant pairwíse correlations (Table S3). The network had low degree (average 4.3 per node), a maximum path of 14, and low checkerboard (C-score= 0.387). The C-score was compared to the null model and found to be different (null model C-score = 0.338) with a 97.5% CI and p<0.001 supporting non-random OTU co-occurrence patterns and the presence of a microbial network. However the work by Berry and Widder (2014) examining the effect of filtering on sensitivity indicates that type II errors may be as high as 0.5 based on 16-20% of the network potentially being habitat specialists (Table S4). Analysis and visualisation of this network (Fig 5) revealed a community consisting of 8 modules.

**Fig 5.**
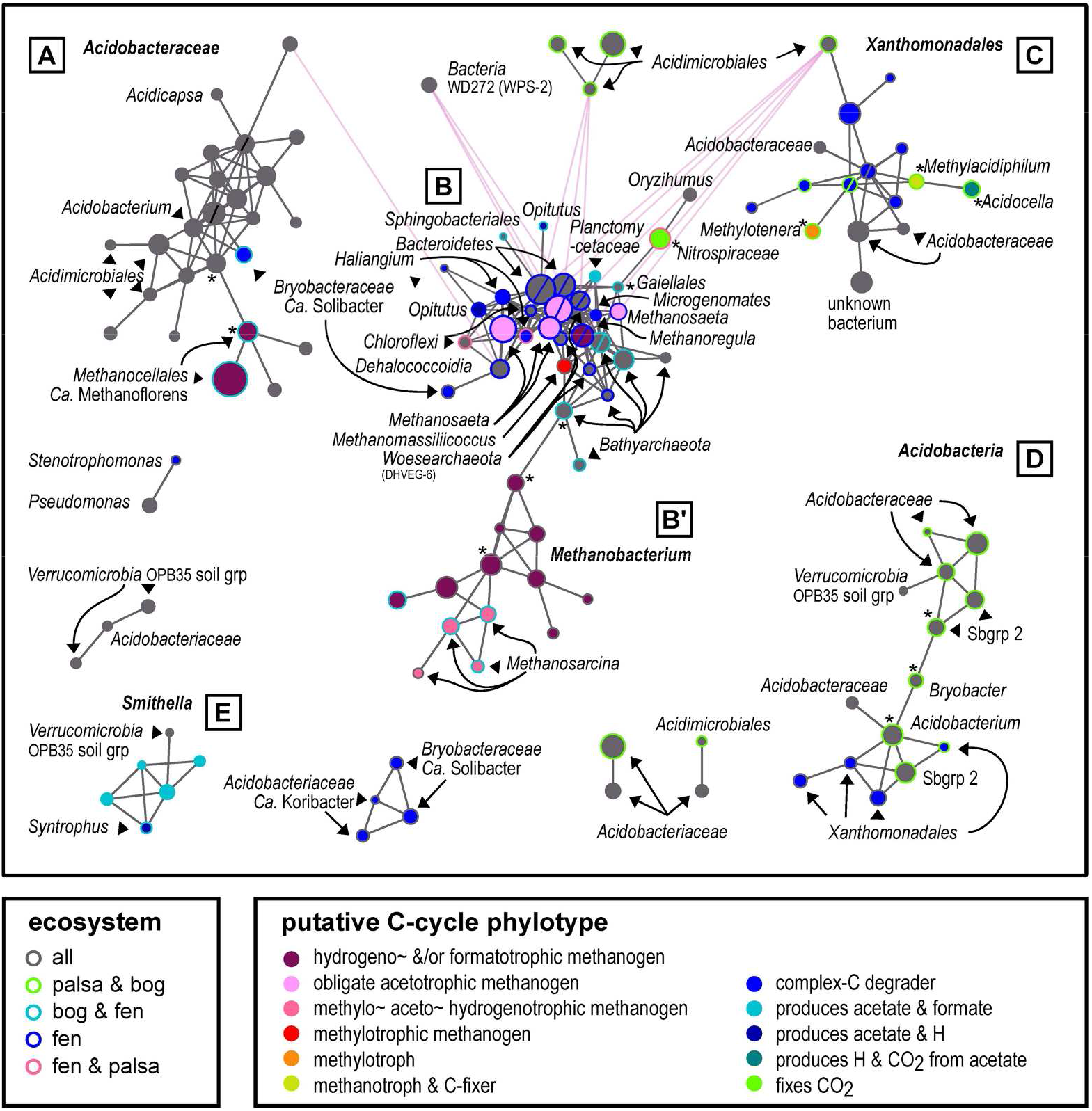
Network of phylotypes present in at least 15 samples with significant pairwise correlation. Circles indicate individual OTUs; circle diameter is a function of the log of the mean abundance. Circle colour denotes phylogenetically-inferred metabolism, and the colour of the outline denotes ecosystem presence. Circles with * are putative keystones and those with a / across the middle are putative hubs. Grey edges indicate co-occurrence, while pink edges indicate mutual-exclusion. 181x184mm (300 x 300 DPI)

The detection of phylotypes in different environments, and their network topological description, enables exploration of community metabolic roles of microbial lineages (Foster et al., 2008). Highly connected phylotypes sometimes called hubs or keystones (high degree, high closeness, low betweenness) are predicted to perform key metabolic steps within microbial communities. In the identified network at Stordalen only hubs with high degree, high closeness but with high betweenness were identified (Fig 5, Table S3 & S4). Due to their high betweenness and close phylogenetic relatedness to adjacent OTUs these hubs exhibit qualities associated with redundancy or ‘niche overlap’ and therefore may have little effect if removed i.e. they are unlikely keystone species. Keystone species, while sometimes described statistically as hubs (Faust and Raes, 2012; Berry and Widder, 2014), are ecologically those that would cause (disproportionate) disruption to a network if lost. In the larger modules of the network, there were a few phylotypes (putative keystones marked on Fig 5, Table S3, S4) that if removed would fragment the network and/or were the only phylotypes identified as associated with a critical metabolic process. The loss of any of the identified keystone phylotypes from any of the three sites, past or future, could affect significant changes in C, N, S, or Fe cycles at Stordalen. Identification of keystone species is problematic if there is a high degree of type II errors or, as is expected with environmental phylogenetic-amplicon surveys, there is limited information available on their phenotypes. Statistically, the predicted keystones in this network cover the full range of betweenness and closeness scores but none had high degree, supporting that high degree is a poor predictor of ‘keystoneness’ in soil microbial communities.

Module ‘A’ consisted of 25 phylotypes (19 Acidobacteria, 4 Actinobacteria, 2 Euryarchaeota) that were dominant in the bog (Fig 5, Table S4). Two of the *Acidobacteriaceae* (subgroup I Acidobacteria) phylotypes were identified as hubs. The potential keystones (based on topology) were another *Acidobacteraceae* and the less abundant of the two *Ca.* Methanoflorens (RCII) phylotypes. *Ca.* Methanoflorens is a hydrogeno/formatotrophic methanogen of the *Methanocellales* order that prefer low H^2^ concentration, and are oxygen and acid tolerant (Sakai et al., 2010; Lü and Lu, 2012; Mondav et al., 2014; Lyu and Lu, 2015). One of the three *Acidobacteria* identified at genus level was a phylotype of *Ca.* Solibacter. Solibacter are capable of degrading complex-C molecules such as cellulose, hemicellulose, pectin, chitin, and starch (Ward et al., 2009; Pankratov et al., 2012), a useful phenotype in the sphagnum-peat of the bog. Another was an *Acidobacterium* that may be able to reduce ferric iron (Coupland and Johnson, 2008). The last of the genus level identified was an *Acidcapsa* which likely preferentially utilise bi-products of sphagnum degradation such as xylose or cellobiose, but if necessary could directly degrade starch and pectin (Kulichevskaya et al., 2012; Matsuo et al., 2016). The four *Acidimicrobiales* identified might contribute to Fe-cycling and likely capable of degrading complex polymers (Kämpfer, 2010; Stackebrandt, 2014). The remainder of module A (unknown *Acidobacteraceae)* are likely chemoheterotrophs that either degrade sphagnum derived polymers or their hydrolysed bi-products (Campbell, 2014). Eighty percent of module A phylotypes had significant positive correlations (ρ ≥ ±0.60, p_ad_<0.001, Table S5) with porewater N, DOC concentration.

Module B ^total^ was the most phylogenetically and phenotypically mixed module, it was further divided into two sub-modules: B and B’. Thirty-four of module B phylotypes were only detected in the fen, eight in both bog and fen samples, and three in both fen and palsa samples. Module B hubs included both of the *Bacteroidetes* phylotypes, a *Woesearchaeota* (DHVEG-6) and a *Methanosaeta,* and a *Methanoregula* and a *Bathyarchaeota* (Msc. Crenarcheaota Grp) (Fig 5, Table S4). The *Bacteroidetes* phylotypes are likely anaerobic, organotrophs with a preference for sugar molecules (Krieg et al., 2010). *Methanosaeta* are obligate acetotrophic methanogens and their higher substrate (acetate) affinity may be one mechanism enabling them to compete in the fen (Westermann et al., 1989; Ferry, 2010; Liu et al., 2010) as would the higher pH and reduction in inhibitory phenolics compared to conditions in the bog. A *Methanoregula* phylotype was also identified as a hub, the only potentially hydrogenotrophic (&/or formatotrophic) methanogen in module B. Its requirement for and greater tolerance of acetate, might assist *Methanosaeta* by using up some of the available acetate (Smith and Ingram-Smith, 2007; Bräuer et al., 2011; Oren, 2014b). No phenotypic data is yet available for *Woesearchaeota.* and the *Bathyarchaeota* so far described were either methanogenic, methanotrophic, or organoheterotrophic (Butterfield et al., 2016; Lazar et al., 2016). So while no predictions can be made as to whether these phylotypes are e.g. methanogens, it is evident that they are important methanogenic-community members. One of the methanogenic phylotypes connected to the Woesearchaeota and Bathyarchaeota was a methylotrophic-methanogen of the *Methanomassiliicoccus* genus (Borrel et al., 2013). Module B and B’ were connected by the co-occurrence of one of these *Bathyarchaeota* and a *Methanobacterium* respectively, both possible keystones. The other putative keystones were an actinobacterial phylotype of the *Gaiellales* order of unknown but likely heterotrophic metabolism, and the chemolithoautotrophic *Nitrospiraceae* (Daims, 2014). It is probable that the *Nitrospiraceae* phylotype, which was not detected in the bog, contributes to the module by C-fixation (Daims, 2014). Two other module B phylotypes not in the bog were a chemoorganotrophic (putative complex-C degrader) *Myxococcales* genus *Haliangium* (Kim and Liesack, 2015), and an uncultured *Chloroflexi* KD4-96. The thirteen OTUs of module B’ were either not detected or detected at very low abundance in the palsa (Fig 5, Table S4). All B’ phylotypes were methanogens from either the hydrogeno/formatotrophic *Methanobacterium* genus or the metabolically flexible *Methanosarcina* genus (Oren, 2014a, 2014c). Module B’, and to a lesser extent B, displays the functional redundancy and phylogenetic clustering characteristic of soil communities and in particular methanogenic soils (Embree et al., 2014). Module B phylotypes (87 %) were strongly correlated (ρ ≥ ±0.60, p_adj_<0.001, Table S5) to phylogenetic even dispersal (NRI), pH, CH4 flux, and decreasing porewater N. Module C consisted mainly of *Xanthomonadales* an order of obligate aerobic *Gammaproteobacteria* capable of degrading complex organic molecules and participating in methyl / H syntrophic partnerships with methylotrophs (Kim and Liesack, 2015). The co-presence of the keystone methylotrophic proteobacterial *Methylotenera* (Doronina et al., 2014) phylotype supports the possibility that such partnerships occur at Stordalen and are common in the aerobic partition of methanogenic soils. A second keystone phylotype was the C-fixing verrucomicrobial methanotroph *Methylacidiphilum* (Hedlund, 2010; Sharp et al., 2013). The final keystone was the putative ferric iron reducing, H_2_/CO_2_ producing *Acidocella.* which may also have a syntrophic partner within module C (Coupland and Johnson, 2008; Johnson and Hallberg, 2008). Most phylotypes (12 of 16) were correlated to increased clustering (NRI) and acidity (Table S5).

Module D consisted of 14 OTUs, nine *Acidobacteria,* four *Gammaproteobacteria (Xanthomonadales),* and one *Verrucomicrobia* all of whom were dominant in the palsa. Three keystones were identified, a subdivision 2 Acidobacteria, a Bryobacter (subdivision 3 Acidobacteria), and a putatively ferric iron reducing Acidobacterium (subdivision 1 Acidobacteria) phylotype. Bryobacter are chemoheterotrophs with a preference for sugars (Dedysh et al., 2016). This module appears to have members that together degrade complex-C molecules, polysaccharides and simple sugars (Hedlund, 2010; Dedysh et al., 2016; Yang et al., 2016). Twelve of the phylotypes had significant correlation (ρ ≥ ±0.60, p_adj_<0.001, Table S5) with distance above frozen ground/watertable. Module E had five *Syntrophobacterales* phylotypes, putative producers of substrates for methanogenesis (acetate, H, and formate) and capable of N fixation (Embree et al., 2014; Lin et al., 2014), with four from the *Smithella* genus and one from *Syntrophus.* None of these five were detected in palsa samples. The final OTU of the module was detected across the mire and identified as belonging to the *Verrucomicrobia* OPB35 soil group which is thought to degrade polysaccharides (Hedlund, 2010; Yang et al., 2016). All of module E phylotypes were significantly correlated with increasing CH_4_ flux and decreasing NRI (ρ ≥ ±0.60, p_adj_<0.001, Table S5).

Overall, the clustering of closely related phylotypes in modules is consistent with the phylogenetic diversity results. The network associations of methanogen phylotypes were complex and modular, with most falling into sub-network B’ (present in bog and fen and low abundance in palsa), followed by B (fen only, except for the more cosmopolitan *Methanomasilliicoccus* phylotype). The two outlying *Ca.* Methanoflorens methanogens in module A (fen and bog) were associated with Acidobacteria and Actinobacteria which are typical of peat environments and The presence of permafrost fits the known distribution of this genus (Mondav et al., 2014) and its predicted phenotype. The putative C-fixing autotrophs were scattered through the network and only moderately connected, supporting their phylogenetically-based assignments to a primary trophic role.

### <Fig 5, 169 mm wide>

#### Known C-cycling phylotypes

Distinct shifts in relative abundances of putative methanogens, and methylotrophs were evident across the Mire. Relative abundance of methanogens increased across the thaw gradient (palsa<bog<fen, Table S6) (K-Wmc, p <0.001). Methanogens were strongly associated with the deepest bog and fen samples (Table S6, Fig S7). All but one of the obligate acetotrophic methanogens *(Methanosaeta)* were detected exclusively in the fen (Fig S7), the other one was found in a single palsa sample. Apart from the anomalous *Methanosaeta* detected in the palsa, this is consistent with reported acetotrophic sensitivity to low pH due to reduction of the ΔG (Gibbs free energy) of the acetotrophic methanogenesis pathway (Kotsyurbenko et al., 2007). The possibility of a divergent metabolism may explain the presence of the *Methanosaeta* in the palsa. Other methanogenic phylotypes detected in apparently aerobic samples (above the water line) may have been enabled by micro-anaerobic-habitats, oxidative resistance as seen in some *Methanocellales* (Angel et al., 2011), or association with an anaerobic host gut (Paul et al., 2012). Putative methano/methylotrophic phylotypes were distributed across all samples (Fig S7) and were highest in the bog samples (K-Wmc, p <0.01,Table S6) likely accounting for the lower CH4 flux despite the abundance of methanogens. Some methylotrophs were detected below the waterline in bog and fen samples (Fig S7) and likely exist in micro-aerobic spaces enabled by plant root gas transport (Colmer, 2003). The relative ratio of methanogen to methanotroph phylotypes differed significantly between sites (K-Wmc, p <0.01), increasing across the thaw gradient (palsa<bog<fen, Table S6). Due to the polyphyletic distribution of autotrophic and methanotrophic metabolisms the assignment of function requires identification to family or genera level, while most methanogens can be identified at class or order level. It is therefore likely that abundances and richness of autotrophic and methanotrophic microbes described here are underestimated more than methanogens. The shifting C-cycling phylotype patterns described here, especially the methanogen to methanotroph ratio provide detail of biogenic methane production and consumption that support reported site C-budgets (Bäckstrand et al., 2010), CH_4_ emissions and CH_4_ isotope ratios (McCalley et al., 2014) from Stordalen.

The presence of permafrost maintains the ombrotrophic, acidic environment (Natali et al., 2011; Tveit et al., 2013) which correlated with a diverse, rich, phylogenetically-clustered, and autotroph-abundant palsa community. The bog sites (thawing permafrost) had significantly lower richness, diversity, evenness and estimated population size than the palsa, the fen, and also other peat bog sites (Lin et al., 2012; Serkebaeva et al., 2013; Tveit et al., 2013). Collectively, this suggests a structural response to ecosystem disturbance (Degens et al., 2001) caused by site inundation as a consequence of subsidence caused by permafrost thaw (Rydén et al., 1980; Johansson and Åkerman, 2008). Correlations between diversity estimates (alpha, beta, and phylogenetic) and distance of sample above or below the water table support that site inundation (and therefore thaw) is a mechanistic driver of community structure and function and that deterministic processes were the main drivers of community composition and assembly in this and other bogs (Quiroga et al., 2015). Complete loss of permafrost in the fen was correlated to assemblages with highest richness, alpha diversity, beta diversity, and phylogenetic even-dispersion.

As the permafrost thaws, causing subsidence, Palsas and transitory bogs at Stordalen mire are expected to give way to fens. At Stordalen the transition from bog to fen is accompanied by community diversification and proliferation of methanogens and a decrease in the relative ratio of methanotrophs. It appears at Stordalen, as found elsewhere under similar conditions (Liebner et al., 2015), that at the point of inundation a regime shift is initiated and that beyond this point, the community does not recover but instead shifts towards a new stable state as found in the fen. The fen assemblage has qualities indicative of greater stability (high redundancy, evenness, richness, and diversity) coincident with an altered C-budget of dramatically higher warming potential. The combination of changes predicted by climate models, the trajectory suggested in biogeochemical and vegetation surveys of the Mire, and the details of the microbial community C-cycling shifts detailed here suggest that the mires along the Torneträsk valley will increasingly add to radiative climate forcing via increased CH_4_ flux over the coming decades as more land is converted to fen. Longer-term outcomes of climate change in this region are projected to eventually include some drying (e.g. terrestrialisation, the conversion of wetlands to drier areas (Payette et al., 2004)) and expansion of the dwarf forests (Rundqvist et al., 2011). If these fundamental habitat shifts occur, they are extremely likely to drive further changes in the microbial communities and C budgets of the region.

### Experimental Procedures

#### Field site and sampling

Samples were taken from the active layer of an individual palsa thaw sequence in Stordalen Mire, subarctic Sweden (68.35N, 19.04E), with three stages of permafrost degradation evidenced by topographical and vegetative characteristics (intact, thawing, and thawed; Fig S1). The intact permafrost was represented by a raised section of the palsa (palsa site, Fig S2); the thawing transition site was an elevationally depressed region within the palsa (bog site); and the thawed permafrost was a thermokarst feature with no detectable permafrost and thus no apparent seasonal active layer (fen site) (Rydén et al., 1980; Malmer et al., 2005). The palsa site was an ombrotrophic, drained, raised peat (altitude 351 m.a.s.l (Jackowicz-Korczyñski et al., 2010)) with tundra vegetation including *Betula nana* and *Empetrum hermaphroditum,* interspersed with *Eriophorum vaginatum, Rubus chamaemorus,* lichens, and mosses. The bog site was a wet ombrotrophic depression sunken ~1 m below the palsa. Vegetation was predominately *Sphagnum spp.* with *E. vaginatum.* Water table varied seasonally from 5 cm above to 30 cm below the vegetation surface and was perched above the local groundwater. The average pH of the ombrotrophic sites was 4.2 +/-0.3 sd. The fen site was a minerotrophic, waterlogged fen ~2 m below the palsa, vegetation was dominated by *Eriophorum angustifolium* with some *Sphagnum spp.* and *Equisetum spp.,* and open water was present. The water table was always within 5 cm of the peat surface, sometimes being above it, and average pH of 5.7 +/-0.1 sd.

Soil cores were taken in August/September 2010 and June, July, August, and October 2011 (Table S1, Table S7). On each of the five sampling dates, between two and four cores were taken from each of the three sites (Fig S1). In 2010 four cores were taken from palsa and bog sites, in October 2011 two cores were taken from the fen site, and all remaining sampling dates and locations had three cores sampled. Samples cut from cores taken from the same site, at either the same depth in cm or the same ecologically significant depth (e.g. depth relative to water table) were designated technical replicates. Samples taken at different depths were analysed as treatments (Samples taken at different depths were selected based on ecologically pre-determined indicators such as at the water table for the bog, Fig S2). Porewater, peat, flux and isotope measurements taken simultaneously to the microbial samples were described in Mondav et al. (2014), Hodgkins et al. (2014) and McCalley et al (2014) but are detailed again in the Supplementary Methods.

#### SSU rRNA gene amplicon sequencing and analysis

Microbial community was surveyed by small subunit (SSU) rRNA gene amplicon sequencing (Mondav et al., 2014). Briefly, DNA was amplified with tagged primers for V6-V8 region of the SSU rRNA gene with the 926F (AAACTYAAAKGAATTGRCGG) and 1392wR (ACGGGCGGTGWGTRC) primers, in duplicate reactions, pooled, and sequenced with the 454 Ti GS (LifeSciences, Carlsbad). Sequences are available from SRA under accession SRA096214 (McCalley et al., 2014; Mondav et al., 2014), SRR numbers and primer details are listed in detail in Table S1. Sequences were cleaned (pre-processed) with MacQIIME v1.9.1 then analysed at an operational taxonomic unit (OTU) of 97% identity. A detailed description of pre-processing methods can be found in the Supplementary Methods. Cleaned sequences were assigned taxonomy using the open picking method and the SILVA database and a phylogenetic tree made with Fasttree2 and manually rooted between the archaeal and bacterial domains (Caporaso, Bittinger, et al., 2010; Caporaso, Kuczynski, et al., 2010; Edgar, 2010; Price et al., 2010; Huson and Scornavacca, 2012; McDonald et al., 2012; Quast et al., 2013). Phylotype lineage obtained by assignment of reads to taxon identity was utilised to assign putative C-metabolism. Phylotypes that were assigned to lineages with known methanogen, methanotroph/methylotroph, and C-fixing members were manually checked before C-metabolism was assigned.

#### Microbial assemblages

Phylum level analysis was obtained by collapsing the normalised OTU table in Qiime, and phyla detected in more than one sample and also present at over 1% in at least one sample were graphed in MS Excel. To investigate dissimilarity of sample and site assemblages a non-parametric ordination (NMDS) was done in R v3.3.1 (R-Core-team, 2011) using the RStudio v0.99.903 (RStudio, 2012) IDE with the vegan package v2.4-1 (Oksanen et al., 2013) and plotted using gplots v3.0.1 and with scales v0.2.3 (Warnes et al., 2011). Environmental variables and parameters were fitted to the NMDS and factors with significant (p < 0.05) correlation plotted. Alpha diversity metrics were generated in Qiime (richness, singletons, Shannon diversity, Fisher alpha, Heip’s evenness, Simpson’s dominance and Chao1 estimates) and analysed in R using the non-parametric Kruskal-Wallis (K-W) followed by post-hoc testing with Kruskal-Wallis multiple comparison (K-Wmc) testing using R package pgirmess v1.6.4 (Giraudoux, 2012). Correlation analyses were done using non-parametric Spearman and linear regression and differences between sites were checked for significance with the largest p value obtained reported. Images were processed for publication in Inkscape 0.91. Analysis of the phylogenetic diversity and distance (PD, NRI, NTI) were calculated using the distance tree output from QIIME, and correlation and equations calculated in R with picante v1.6-0 (Gotelli, 2000; Faith, 2006; Kembel et al., 2010). OTUs present in less than 15 samples were removed and the resultant OTU table analysed for pairwise interactions in MENA (Deng et al., 2012), fastLSA (Durno et al., 2013), CoNet v1.1.1 (Faust et al., 2012) and SparCC (Friedman and Alm, 2012). All networks were loaded into R and OTU pairs that were identified as present in at least two of the four networks were retained and the network analysed in CytoScape v3.4.0. See Supplementary Methods for detailed description of all methods.

Supplementary Information is available as a separate download and includes Supplementary Methods, Figs S1-S7, Equations S1-S5, and Tables S1-S7.

## Acknowledgements

Many thanks to Andrew C Barnes, Brian Lanoil, and James Prosser for critical comments on a previous version of the manuscript. Sincere thanks to the two anonymous reviewers who helped me greatly improve this manuscript. RM was supported by an Australian Postgraduate Award Scholarship and a Swedish Vetenskapsrådet grant. JPC, PMC, SF, SH, CKM, SRS, and VIR were supported by the US Department of Energy, Office of Biological and Environmental Research under the Genomic Science program (Award DE-SC0004632).

## Author contributions

JPC, PMC, SF, SRS, VIR designed the project. RM designed and performed all bioinformatics analyses and visualizations. RM wrote the paper in consultation with all authors.

